# Incredible internal strains within a biogenic single crystal viewed by X-ray diffraction tomography

**DOI:** 10.1101/2020.11.16.384610

**Authors:** Eva Seknazi, Paul Zaslansky, Alex Katsman, Julie Villanova, Boaz Pokroy

## Abstract

The dorsal arm plates (DAPs) of the *Ophiocoma Wendtii* brittle star are highly functional single crystalline biominerals whose optimized structure and nanostructure enable them to fullfill mechanical and optical functions in the organism. Here, a large DAP bulk piece is characterized by means of synchrotron X-ray Diffraction Tomography (XRDT). This non-destructive crystallographic characterization revealed an astounding feature: the presence of very high compressive strains which relax when the mineral is cracked or grinded into a powder. Thus, previous destructive characterization techniques did not allow their detection. We attribute the compressive strains to the previously identified high-Mg calcite particles, which are coherently included and thereby compress the low-Mg calcite matrix. The measured slice contained both the bulk DAP sample as well as DAP powder. The data generated by the bulk piece could be separated from those by the powder, and the latter was used to calibrate and interprete the former. This study reveals yet another awe-inspiring feature of a biogenic structure, highlights the importance of non-destructive crystallographic characterization for biominerals, and exemplifies the potential of XRDT use in studying a single crystalline material, as well as the advantage of complementary measurement of bulk and powder for data calibration and interpretation.

## Introduction

Biominerals are the minerals produced by living organisms. They are riveting materials that often display intricate structures, associated with properties far superior to those of their geological counterparts.^1–4^ Their structures are often fascinating, complex, and bear Nature’s design strategies for highly functional materials.^5–9^ Studying them can lead to valuable lessons for the design of new materials.^10–15^ Given the multiplex and hierarchichal architecture of biomenrals over multiple lengthscales, investigation of their internal structure with non-destructive characterization techniques is essential. Accordingly, non-destructive tomography techniques, such as micro-computed tomography or holotomography have been used to obtain 3D imaging of the interior structure of biominerals.^16–19^ We employ here a lesser-used XRDT technique. It allows to acquire 2D diffraction data at several positions and orientations of the sample and, via tomographic reconstructions, to map the regions in the sample at the origin of observed features in the diffraction data. XRDT was previoulsy used on polycrystalline materials, whose 2D diffraction data display characteristic ring patterns.^20–24^ In those studies, crystalline phases and texture could be spatially resolved by reconstructing full rings or ring sections from the acquired data.^20–24^ However, in the case of single crystals the acquired diffraction patterns are not ring patterns but rather isolated diffraction spots, wich appear at specific orientations of the specimen. To the best of our knowledge, no study involving XRDT analysis of a single crystal has been carried out. We apply here XRDT on a large biogenic single crystal: the dorsal arm plates (DAPs) of the *O. wendtii* brittle star.

The DAPs of the *O. wendtii* brittle star demonstarte an exceptionnally level of crystal engineering and tailored functional. They are single crystals of calcite, a brittle and birefringent mineral, and yet serve the mechanical and optical purposes of the organism.^25–28^ This is made possible by their tailored structure, that confers them toughness and optical capabilities.^26–30^ Indeed, the DAPs contain hundreds of rounded features, called lenses, whose rounded shape guides the light through and focuses it onto photoreceptor nerve bundles. In addition, lenses’s alignment along the optical *c*-axis of calcite minimizes birefringence.^26^ We have recently showed that the lenses’ toughness is more than twice the toughness of geological calcite, owing to an elaborate toughening strategy based on a specific distribution of Mg, that substitutes for Ca in the calcite lattice.^27^ The total content of Mg in the DAPs calcite is 15% (*i.e.* Mg/(Ca+Mg)=0.15) but the distribution of Mg is inhomogeneous whereby the single crystals are comprised of two coherent Mg-calcite phases: a matrix of low-Mg-calcite, and coherently included high-Mg calcite nanoparticles.^27^ Given that high-Mg calcite lattice is smaller than that of low-Mg calcite, as Mg ions are smaller than Ca ions,^31^ the coherency between the two phases induces tensile strains in the high-Mg particles and compressive strains in the matrix.^27^ Moreover, the high-Mg nanoparticles were shown to be distributed in alternating particles-rich and particles-depleted layers, so the compressive strains in the material are in fact layered.^27,29^ The existence of coherent high-Mg nanoparticles and associated strains are responsible for the enhanced mechanical properties of the otherwise brittle lenses.^27,30^ The crystallographic characterization of atomic structure via synchrotron High-resolution Powder X-ray Diffraction (HRPXRD) of powdered samples: the diffraction pattern displayed peaks correponding to a single low-Mg calcite phase. Upon heating, the coherent high-Mg particles grow within the lenses and lose coherency with the matrix, thereby incoherent high-Mg particles’ diffraction peaks become visible by HRPXRD.^27^

HRPXRD allows the extraction of structural parameters with high precision but contains a drawback: the samples must be finely powdered. In contrast, XRDT provides crystallographic data of a large bulk sample, undamaged by sample preparation steps. We show that XRDT data of a DAP sample reveal structural features that were undetectable from the analysis performed on the powdered sample: the existence of high compressive strains, which relax when mechanichally damaged or heated. Such finding, among others, was enabled by the combination of studying powdered DAP combined with the large whole DAP measured piece, that was simultaneously measured and could be used for calibration and data interpretation.

## Results and discussion

An *O. wendtii* DAP piece was measured by both 3D Nano-holotomography (Nano-HT) and 2D XRDT. The selected plane for the XRDT measurement was reconstituted by reconstructing the Nano-HT data and corresponded to a lens cross-section (Figure 1a). The position of the 2D slice measured with XRDT within the 3D reconstructed volume is shown in Supplementary Figure 1. In this XRDT experiment, the sample rotated, until a 180° rotation was completed; at each angle, horizontal line scans were performed. A diffraction pattern was acquired at each of the positions, at each angle of rotation. The data set therefore consists of a frame stack, each frame is a diffraction pattern, the total number of frames is equal to the number of angles multiplied by the number of translations per angles.

**Figure 1.**
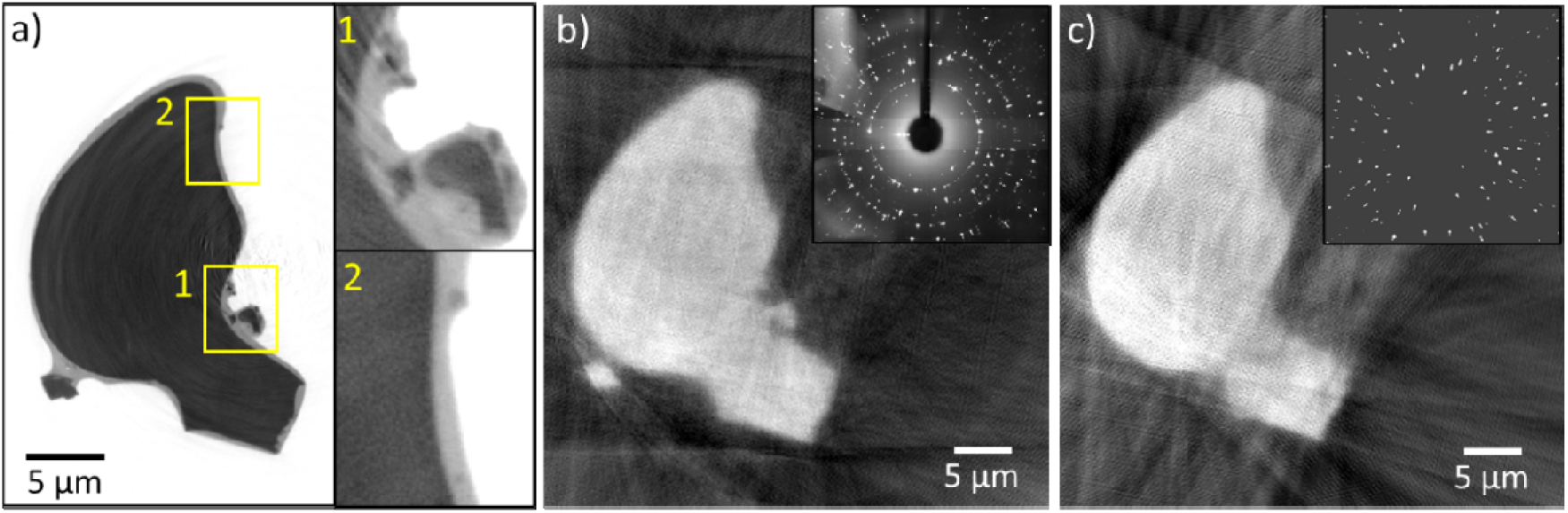
a) Nano-HT reconstructed 2D slice of the plane measured by XRDT (pixel size 25 nm), insets: debris near the lens, b) reconstruction of the total intensity of the diffraction patterns, inset: average frame c) reconstruction of the total intensity of the filtered data showing the removal of debris, inset: average frame of the filtered data.

The brittle star DAPs are single crystals of Mg-calcite. Because of their single crystalline nature, a reflection corresponding to one set of crystallographic planes appears at a specific rotation angle only, when the crystallographic planes are parallel to the beam. Correspondingly, a single frame among the full data set displays a few reflections only (Supplementary Figure 2a). In order to analyze the different reflections simultaneously, we considered an average frame. We created this average frame by creating the stack of maximum intensity projection of the frames acquired at the same rotation angle and averaging it. The average frame of the whole data set therefore displays all the reflections (Supplementary Figure 2b). Tomographic reconstructions enable to locate the regions in the sample that are responsible for the intensity of selected pixels in the diffraction pattern frames. It is therefore possible to select pixels (the selected pixels constitute the region of interest (ROI)) in the average frame and measure the intensity of those pixels in all the acquired diffraction patterns, and finally to reconstruct the intensity profile and locate the regions in the sample generating this intensity.

XRDT reconstructions of a single crystal are however intricate, as each reflection seen in the average diffraction pattern frame appears at one specific rotation angle only. Therefore, when a selected ROI in the average frame consists of one single reflection, the measured intensity within the ROI is high at a specific angle only. Thus, the spatial information given by reconstruction of one reflection is unidimensional: it is the projected intensity of the reflection along the sample on a specific direction (Supplementary Figure 3). Therefore, in order to spatially locate the areas in the sample responsible for a certain feature observed in the average diffraction pattern frame, at least several reflections should be selected and reconstructed together so that spatial information would be obtained at different angles. By selecting all the pixels of the average frame (*i.e*. the ROI consists of the entire frame) and reconstructing the total intensity of the diffraction patterns, one should obtain the shape of all crystalline objects in the measured plane. Indeed, such reconstruction yielded the shape of the lens as well as the shape of DAP debris near the lens (Figure 1b), that are identical to those from the Nano-HT reconstructed slice shown in Figure 1a.

As the debris are crystalline, they generate reflections that are visible in the average frame. In order to separate the reflections generated by the debris from those generated by the bulk lens in the average frame, we created a filter and applied it to the raw dataset. The filter recognizes the intensities originating from debris using the discriminating characteristic that the debris generate intensity in a small number of frames, as they are small. The filter then replaces the intensities generated by debris or by the background by a constant value (see Supplementary Figure 4 and experimental section). The remaining intensities are those generated by the bulk lens itself. Averaging the filtered data indeed yielded an average frame without background and with much less diffraction spots than in the average frame of the unfiltered data (Figure 1b, c, insets). Moreover, reconstructing the total intensity of the filtered frames (*i.e*. the ROI consists of the entire frame) yielded the shape of the bulk lens, without the debris (Figure 1c). This reconstruction confirms that the filter removed the debris, and that the debris were responsible for the polycrystalline-like diffraction rings in the unfiltered average frame. We note that other than the two micrometer-size debris pieces that can be resolved by the XRDT reconstruction (Figure 1b), the measured slice contains several smaller debris, that can be seen in the Nano-HT reconstructed slice, and are responsible for the polycrystalline-like ring pattern of the unfiltered average frame (Figure 1a, insets).

By analyzing the unfiltered and filtered average frames (Figure 1) and zoom-ins (Figure 2) a few observations can be made. The first observation is that most of the reflections generated by the bulk lens (*i.e*. reflections present in the filtered data) are radially elongated, they are in fact twofold. This indicates that the single-crystal bulk lens actually posseses several strain states within its lattice. The second observation is that the radii of the polycrystalline-like rings caused by the presence of debris are those of the smaller-radii diffraction spots of the twofold reflections from the lens. Since the debris can be assimilated to powdered lens, the radii of the polycrystalline-like rings correspond to the *d*-spacings measured by powder XRD. Accordingly, the additional diffraction spots at larger radii correspond to *d*-spacings that are smaller than those measured by powder XRD. These diffraction spots therefore attest for a compressive strained state of the lens lattice, with smaller lattice parameters than those found with powder XRD. The smaller-radii diffraction spots, corresponding to the lattice constants found by powder XRD, are referred to as the unstrained state of the lens lattice.

**Figure 2.**
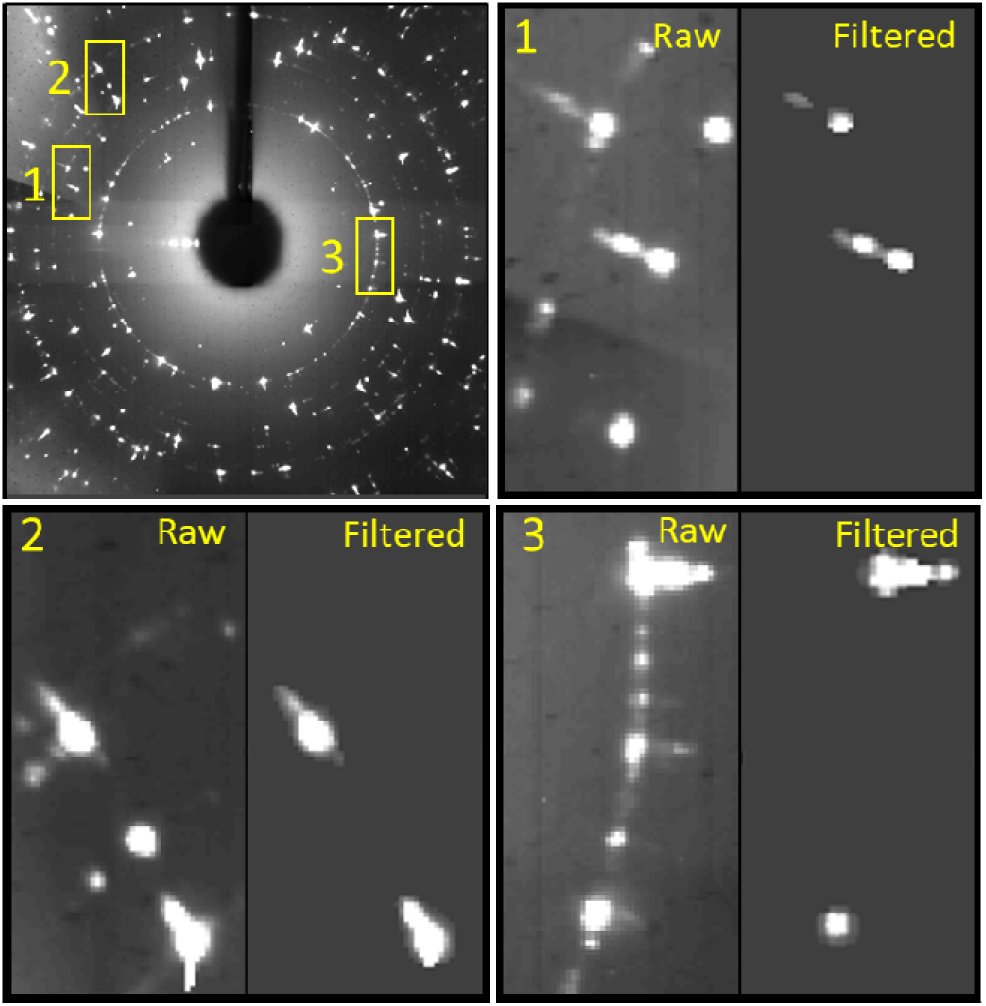
Average frame and zoom-ins of the unfiltered and filtered average frames in three different regions.

By separately plotting the intensities of pixels associated with the unstrained and strained parts of reflections through the dataset frames, one can notice a certain characteristic behavior. Whereas the intensity of pixels associated with the strained part of the reflection is smooth (through the frames acquired at the relevant rotation angle), the intensity of the pixels associated with the unstrained part is spiky (Supplementary Figure 5). The latter indicates that there are localized regions in the lens that are responsible for very high intensity of the unstrained reflections. However, apart for those spikes, the intensity of the unstrained reflections is not necessarily higher than the intensity of the strained reflections, as can be seen in Supplementary Figure 6 and Supplementary Figure 6.

In order to locate the regions in the lens responsible for the very high intensity of the unstrained reflections, we reconstructed separately the intensity of the unstrained and strained parts of several reflections. We used the filtered dataset so the reconstructed intensities will be of the bulk lens only, and without background. We used 44 different double reflections; although there is no intensity at all the rotation angles, it is a sufficient amount to recognize the lens shape, in both the reconstructions of the intensities of the unstrained and strained parts of the reflections (Figure 3a, b). The reconstruction of the unstrained parts of the selected reflections indeed displays localized areas in the sample with much higher intensity (Figure 3b). On the contrary, the reconstruction of the strained parts of the selected reflections displays smooth intensity throughout the lens (Figure 3a).

**Figure 3.**
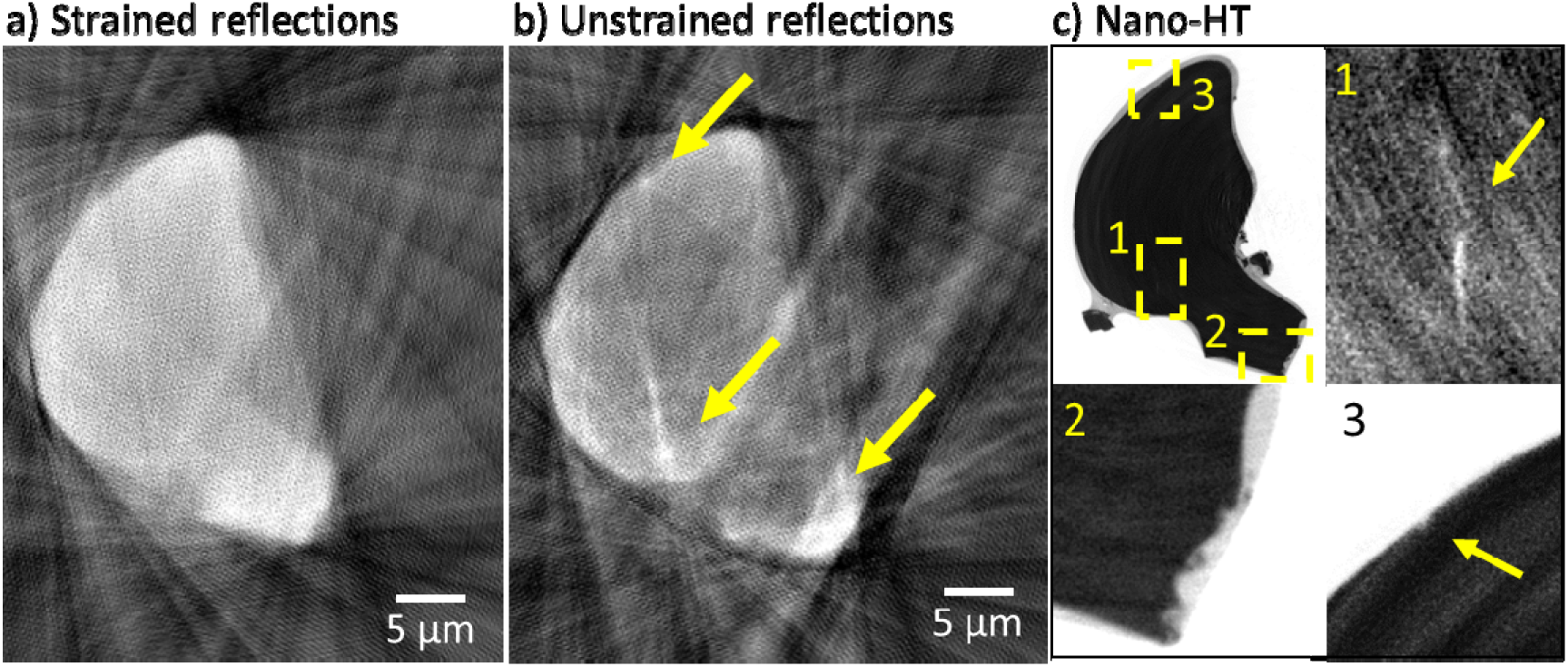
a) Reconstruction of the pixels corresponding to a) the strained and b) the unstrained reflections showing higher intensity in localized areas indicated by yellow arrows. c) Nano-HT slice corresponding to the plane measured by XRDT and zoom-ins of the unstrained regions as localized by XRDT.

The surprisingly large intensity differences over confined regions in the reconstruction of the unstrained reflections’ intensity was elucidated by the examination of the high-resolution Nano-HT slice (Figure 3b). These localized high-intensity areas correspond to the mechanically damaged places: an interior crack or dented surfaces. The interior crack can be better appraised in Nano-HT slices at slightly different heights, where the crack is more open (Supplementary Figure 8). The fact that mechanically damaged places in the lens correspond to much higher intensity of the unstrained reflections indicates that these mechanical damages are associated with significant strain release. The latter is corroborated by the fact that the debris, which endured grinding to be detached from the bulk piece, generate reflections that mainly correspond to the unstrained state (Figure 2). Therefore, the debris and the damaged parts of the lens released considerable amounts of strain, causing localized unstrained state. On the other hand, the strained state is rather uniformly distributed within the bulk single crystal lens (in the resolution limit imposed by the technique). Moreover, when no damaged parts are crossed, the intensity from the strained state’s reflections can largely prevail the intensity from the unstrained state’s reflections (Supplementary Figure 5 and Supplementary Figure 6).

We considered whether the high intensity of the unstrained reflections at defects are due to dynamical scattering effects. The theory of dynamical diffraction is indeed applicable to large single crystals and predicts high contrast at defects, such as cracks.^32^ However, assuming the lens crystal lattice is regular enough for dynamical diffraction to occur, despite the presence of Mg and high-Mg inclusions, the fact that the unstrained and strained reflections evolve independently or on the expense of one another in the sample (Supplementary Figure 5b and c), indicates that dynamical diffraction is not the key for interpreting these data. Moreover it also demonstrates that the high intensity of the unstrained reflections at defects and in case of powdered DAPs correspond to mechanically-induced strain release.

Such strain release probes the high extent of internal elastic strain present in the lenses and is specific to the lens structure. Indeed, no such behavior was seen in a similar XRDT experiment performed on the narrowly distributed large crystals of another calcium carbonate biomineral: the vateritic spicules of *Herdmania momus* (Supplementary Figure 7). Therefore, such behavior is intrinsic to the lens structure, in which elastic strains are known to exist owing to the presence of high-Mg nanoparticles, which are coherently included in a low-Mg matrix.^27^ We note that as the high-Mg nanoparticles are coherent with the low-Mg matrix, and are of small size (~5 nm diameter^27^), they possess lattice parameters that are identical to the matrix. However, the lattice parameters of the matrix are reduced (strained) by the presence of the numerous coherent high-Mg nanoparticles. It is therefore reasonable to advance that the strains caused by the presence of the coherent high-Mg nanoparticles are at the origin of the observed strained reflections in XRDT. Such strains were invisible in powder diffraction as they were released by powdering the sample. As these strains arise from the coherency and lattice mismatch between the high-Mg nanoparticles and the low-Mg matrix, the strain release generated by mechanical damage may be caused by a partial loss of coherency, and a relief of misfit strain. Such misfit strain relief, mediated by the creation of a dislocation, was observed in another calcite system of synthetic calcite, that was epitaxially grown on SAMs.^33^ Another reason of strain release during cracking is relaxation of macrostrains caused by Mg-rich particle concentration gradients present in the layered structure.

In order to quantify the strains associated with the twofold reflections, we calibrated the diffraction pattern, indexed different reflections, and finally converted the radius of selected reflections, in pixels, to units of *d*-spacings and lattice parameter. The calibration of the diffraction pattern was facilitated by the presence of the polycrystalline-like rings in the average frame caused by the debris. Indeed, the lattice parameters associated with the debris’ polycrystalline-rings are known, as they are extracted from powder XRD. Moreover, the radius (in pixel) and center of the bright {104} ring could be easily extracted and then used in order to find the sample-detector length. The radii (in pixel) corresponding to other *d*-spacings could then be calculated. The rings corresponding to the *d*-spacings of other diffracting set of planes were drawn on the average frame so the reflections could be indexed to their corresponding family of planes (Figure 4).

**Figure 4.**
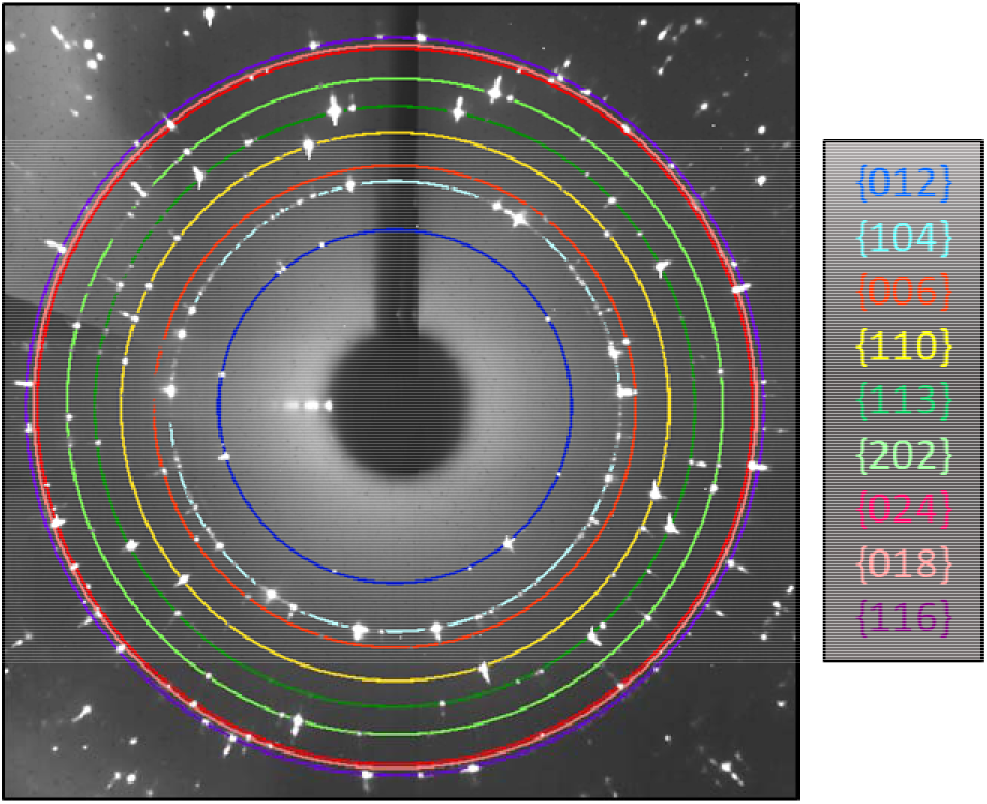
Average frame with depicted colored circles whose radii correspond to the families of calcite diffracting planes.

The most straightforward way to calculate the *a* and *c*-lattice parameters of the strained calcite is to analyze the position of the strained spots of the {110} and {006} reflections respectively, relatively to their unstrained position. We plotted the radial intensity profile of such reflections and extracted the distance between the peaks corresponding to the unstrained and strained states. The {110} and {006} strained reflections were radially shifted of respectively 0.0151 ± 0.0009 Å^−1^ and 0.0109 ± 0.0009 Å^−1^ (Figure 5, Supplementary Figure 9a, b). These radial shifts correspond to the lattice parameters of *a* = 4.755 ± 0.010 Å and *c* = 16.284 ± 0.040 Å. If compared to the lattice parameters of the unstrained state (*i.e.* lattice parameters extracted from powder XRD), these lattice parameters correspond to lattice strains of: and. These calculated lattice parameters indeed fit the position of other reflections’ strained parts (Supplementary Figure 10).

**Figure 5.**
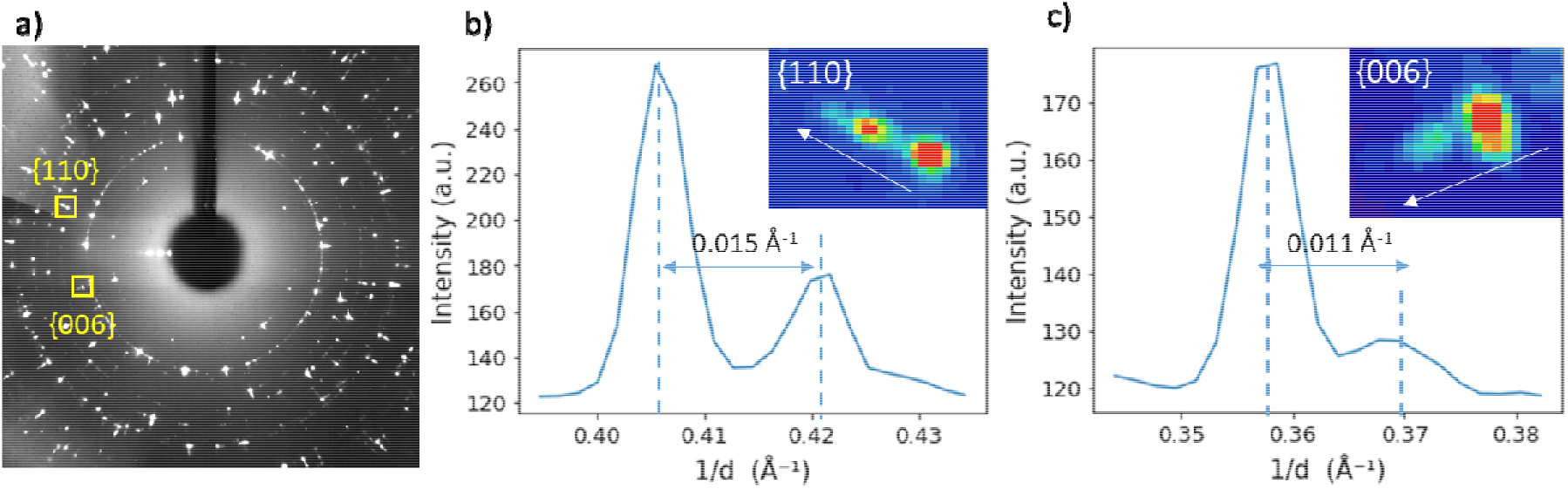
a) Average frame and the reflections considered for strain estimation. b) Radial intensity profile of a {110} reflection showing two distinct peaks. c) Radial intensity profile of a {006} reflection showing two distinct peaks. The insets show the reflections and the arrows show the direction of increasing radius.

The extent of these strains is surprising for two reasons. The first reason is that they are remarkably high. Assuming the discovered compressive strains are solely due to the presence of coherent high-Mg particles, the maximum possible compressive strains in the matrix are equal to the mismatch between the matrix and nanoparticles’ lattices multiplied the volume fraction of the particles, ϕ^25^. If the nanoparticles contain 60% Mg, the lattice mismatch would be of 3.4% and 5.5% for the *a* and *c*-lattice parameter respectively (as calculated using the previously established equations linking the lattice parameter of calcite with the amount of Mg substituting Ca^34^). However, the average compressive strains in the matrix should be much smaller than the lattice mismatch since the average volume fraction of high-Mg nanoparticles is usually less than 10%. Nevertheless, since the elastic constants of high-Mg calcite are higher than those of low-Mg calcite;^35^ the misfit strains are concentrated in the matrix as compressive strains, rather than in the nanoparticles as tensile strains. Therefore, the compressive strains in the matrix should be close to the particles-matrix lattice mismatch.

We note that the exact content of Mg in the high-Mg nanoparticles as well as the volume fraction of these particles are unknown as they could be characterized after heating the DAP powder only, after they grow, lose coherency with the matrix, and become detectable via powder HRXRD.^27^ The content of Mg in the heated, grown and incoherent high-Mg nanoparticles was found to be of ~40% Mg^27^.

It should be noted that lattice parameters of the unstrained state, *a*_un_ = 4.921 ± 0.010 Å and *c*_un_ = 16.675 ± 0.033 Å correspond to average concentration of Mg ~15%. If the Mg-rich particles remain coherent with the low-Mg calcite matrix after cracking, the micro-strains caused by the mismatch between the lattice and particles remain unreleased producing compressive stress in the matrix and tensile stress inside the particles as described in our previous work^25^. Therefore, the discovered strains within the bulk single crystal lens could not be simply the microstrains caused by the average lattice/particle mismatch, but most probably are the result of micron scale gradients of Mg concentration caused by formation of the Mg-particles layered structure. In a such case, the lattice parameters in the region of very high concentration of Mg-rich particles are indeed would be close to ones inside the particles, while the rest of coherent calcite matrix between the high concentration layers would like to have larger lattice parameters, but elastically strained due to required compatibility of the whole lattice structure. If averaged concentration of Mg in the layered structure changes non-linearly between the parallel high-Mg regions (walls), from a maximum value at the wall to a minimum value in the middle between walls, corresponding unstrained lattice parameters changes also non-linearly from some minimum values to maximum values. If the layered structure is parallel to (001) crystallographic plains of calcite, the stresses are caused only by the change in *a* lattice parameter. Assuming constant elastic parameters throughout the layered structure, and some symmetric function for the local volume fraction of the particles, η*(z)*, providing symmetric physical deformations, *f(z)* = (Δ*a*_*p*_ / *a*_0_)η*(z)*, where Δ*a*_*p*_ / *a*_0_ is the relative change of lattice parameter, a, inside the particles, *z* ∈ [−h/2,+h/2], *h* is the layered structure step, one can find elastic strains and stresses between such walls:

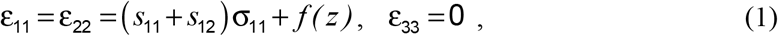

where *s*_11_ and *s*_12_ are the compliance tensor components of calcite, *s*_11_ + *s*_12_ = *C*_33_/*k*, 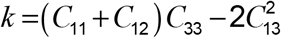 *C*_ij_ are the stiffness tensor components. Applying the Saint-Venant principle stating that resultant force and resultant moment on the contour of the layer are equal to zero, one can obtain:

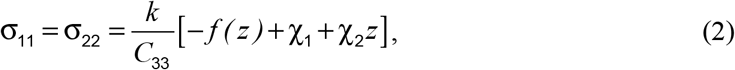

Where 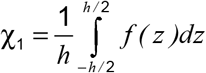 and 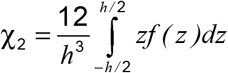. For symmetric function *f(z)* the second integral equals zero, χ_2_ = 0. Therefore, elastic strains in the layer are the following:

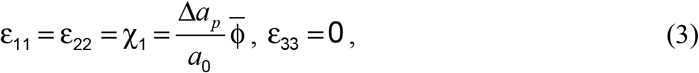

Where 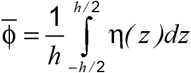 is the average volume fraction of Mg-rich particles over the layered structure. One can see that these strains could not reach so high values as were measured, since the average volume fraction of Mg-rich particles is probably less than 10%. To explain this discrepancy, one can assume that elastic constants also depend on the volume fraction of the particles, and could be much higher in the regions of high particle concentrations. In a such case, deformations in the softer regions (low particle concentrations) can be estimated as

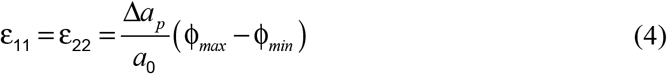

where the difference of maximum and minimum volume fractions, (ϕ_*max*_ ϕ_*min*_), between “walls” and “valleys” of the layered structure may reach the values of (0.65-0.75). Since 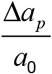 may be of about 0.04 (at η~0.5) one can expect elastic strains ɛ_11_ ~(2.6÷3.0)%, which are comparable with the measured values.

Another point to be understand is the simultaneous compressive elastic strains in ****a**** and ****c**** directions. This can manifest that the layered structure is not parallel to (001) plains. If, for example, the layers are parallel to (104) plains, and the physical strains along (104) plain are ɛ_0_, the corresponding deformations in *a* and *c* directions can be written as:

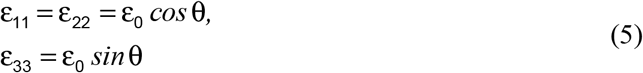

where θ is the angle between (104) and (001) plains, 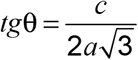 In this case, 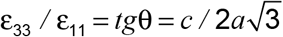. Using the average lattice parameters of unstrained matrix presented above, *a* = 4.921 Å and *c* = 16.756 Å, one can find ɛ_33_ / ɛ_11_ ≈ 0.98. This is comparable with the experimental ratio 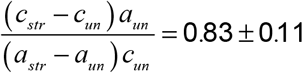. On the other hand, the difference can be connected with the platelet-like shape of the nanoparticles (see below).

The discovered high strains in the untreated DAP sample may indicate that the content of Mg in the unheated high-Mg particles is higher than the one in the particles after heating. Since the high-Mg particles grow with heating at the expense of the low-Mg matrix, it is indeed possible Since the high-Mg particles grow with heating at the expense of the low-Mg matrix, it is indeed consistent that the Mg content of high-Mg particles would decrease as they grow.

We note that we do not consider here another common source of strains in biominerals, namely, the presence of intracrystalline organics.^36^ These are not considered, as they are typically responsible for tensile strains usually of much lower magnitude than those observed here, and were found to be in very low amount in the *O. wendtii* DAPs.^29^

The second reason for finding the discovered strains surprising is that the distortion along the *a*-parameter is larger than that along the *c*-parameter. Typically, distortions in calcite are significantly larger along the *c* than along the *a*-parameter, owing to the smaller elastic constant along the *c*-parameter (*C*_11_/*C*_33_ = 1.8).^37^ The fact that the *a*-lattice parameter of the coherent high-Mg nanoparticles is more strained than the *c*-lattice parameter corroborates the fact that the high-Mg particles in the bulk lens have a platelet-like shape, with the platelet flat face parallel to the lens surface. The latter was indeed inferred via SAXS and HRTEM measurements.^27^ Since the lens surface is oriented along the *c*-axis,^26^ the platelets flat face is parallel to the *a*-direction whereas the thin side is parallel to the *c*-direction. In other words, the effective surface fraction of particles in a-direction, ϕ_a_, is higher than that in c-direction, ϕ_c_; their ratio can be estimated as ϕ*_a_* / ϕ_*c*_ = *l_a_* / *l_c_*, where *l_a_* and *l_c_* are dimensions of platelets in a and c directions. Since in unheated samples *l_a_* / *l_c_* □ 1, the strains along a-axis turned out to be larger than along c-axis.

The *c*-axis of the high-Mg particles therefore withstands higher tensile strains from the coherent low-Mg matrix than its perpendicular plane. Reciprocally, the *a*-axis of the matrix withstands higher compressive strains from the coherent high-Mg particles than its *c*-axis. Therefore, the value of *a*-lattice parameter of the strained material is further from that of the unstrained material, and the distortion is larger.

It is known that heating of this biomineral significantly alters its structure: the coherent high-Mg nanoparticles grow and lose coherency with the matrix, as observed by HRTEM coupled with in situ heating.^27^ Therefore, we expect the strains associated with the coherency and lattice mismatch between the high-Mg nanoparticles and the low-Mg matrix to be relieved upon heating. To verify this, we have performed XRDT on a DAP piece that was heated (400°C, 30 min) prior to the measurements. By performing similar analysis as for the untreated sample, we found that additional diffraction spots at slightly larger radii exist as well, for all reflections, indicating the presence of unrelieved strains. However, unlike the untreated sample, the intensities of the strained reflections in the heated sample were consistently inferior to the unstrained reflections’ intensities (Supplementary Figure 11). Therefore, strain release indeed occurred as a result of heating the mineral. The strained reflections correspond to lattice parameters of *a* = 4.806 ± 0.010 Å and *c* = 16.281 ± 0.040 Å (Figure 6, Supplementary Figure 9c, d). These lattice parameters correspond to lattice distortions (relative to the unstrained lattice parameters of the lenses) of 2.4 ± 0.2 % and 2.9 ± 0.2% in *a* and *c*-lattice parameters, respectively. One can note that although the strains along the *c*-axis are unchanged upon heating, the strains along the *a*-axis are significantly reduced. The latter is probably caused by the partial coherency loss in *a*-direction during growth of high-Mg nanoparticles, from thin platelet-like shape to a less-strained, more equisized shape.

**Figure 6.**
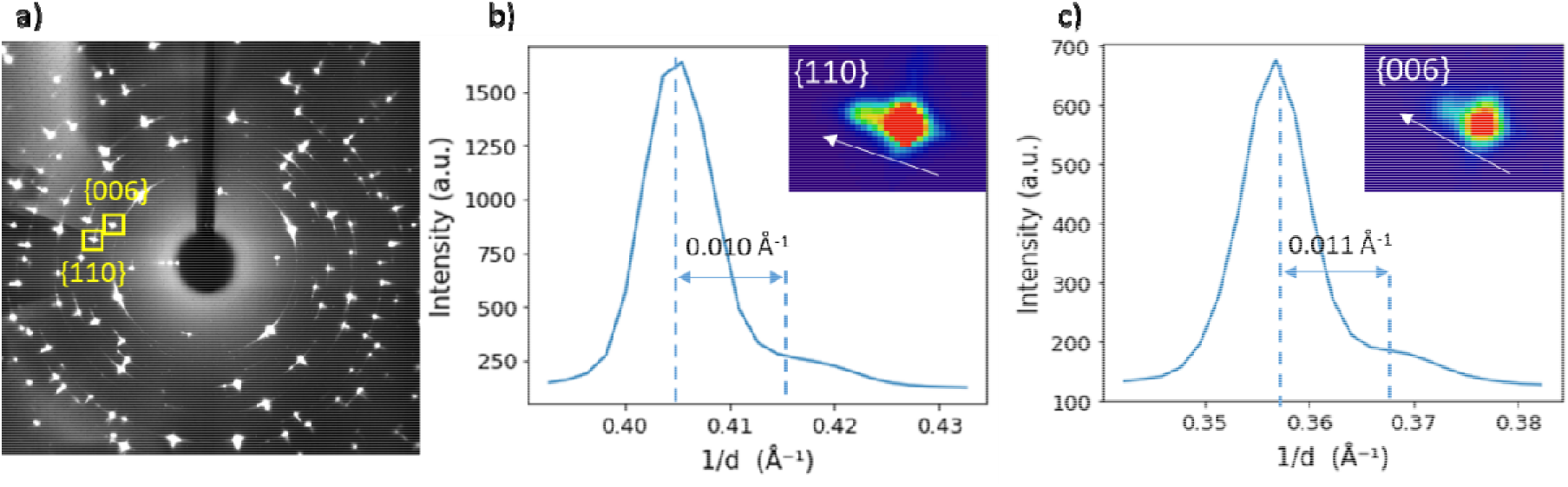
a) Average frame and the reflections from heated sample considered for strain estimation. b) Radial intensity profile of a {110} reflection showing two distinct peaks. c) Radial intensity profile of a {006} reflection showing two distinct peaks. The insets show the reflections and the arrows show the direction of increasing radius.

In addition to the decrease in the intensity and distortions associated with the strained reflections in XRDT, heating the bulk DAP provokes a striking behavior of the biomineral: the lenses burst. The latter could be observed via Nano-HT of a DAP piece before and after heating, where most of rounded lenses become collapsed after heating (Figure 7). The collapse of the rounded lenses is in fact the result of bursting, as can be seen in the radiograph movie acquired while heating a bulk DAP sample (Supplementary Movie 1). Other than the rounded lenses, other parts of the DAP are damaged as a result of heating, as can be seen from the pores visible on a Nano-HT reconstructed slice (Figure 7c). These damages in the integrity of the heated DAP and resulting misoriented regions can explain the azimuthal broadening of the reflections visible in the average XRD frame (Figure 6a). We deduced that the bursting of the lenses is a manifestation of strain and elastic energy release which stems from the coherency loss of the high-Mg nanoparticles. Thus, the high tensile strains inside the particles present in the biomineral are such that their release causes multiple microcracking and breaks the lens.

**Figure 7.**
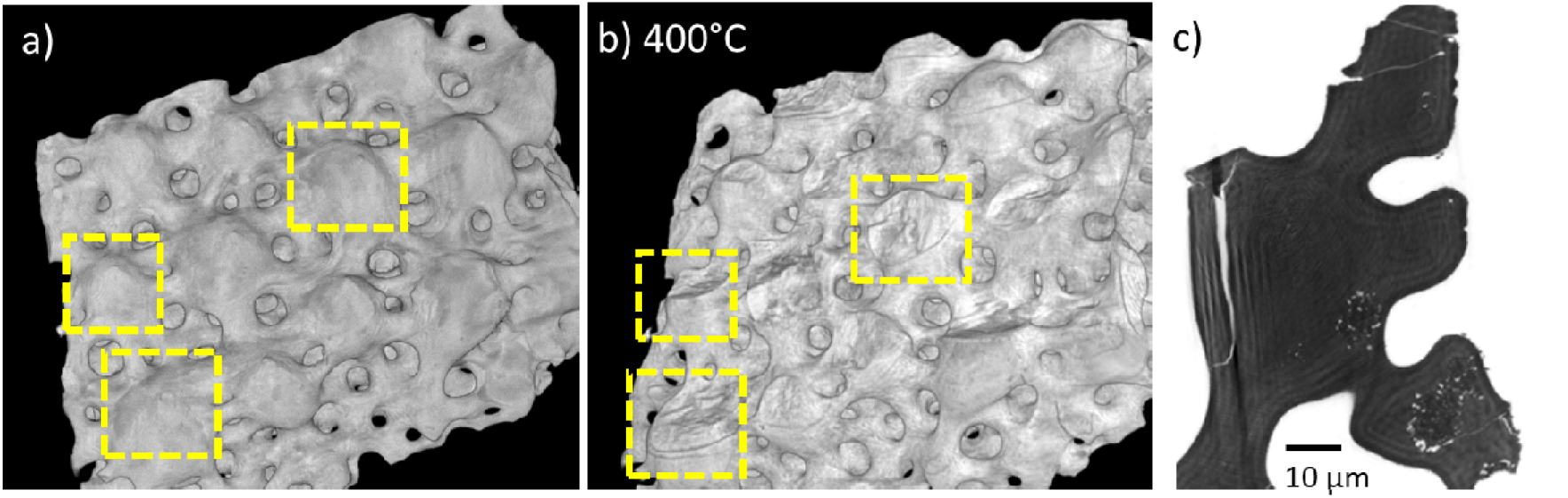
a, b) 3D nano-HT reconstructed volume of a DAP sample at room temperature (a) and after heating at 400°C (b), inside the dotted squares are lenses that burst as a result of the heating (pixel size: 120 nm). c) 2D nano-HT reconstructed slice of a heated DAP sample (pixel size: 50 nm) showing the damage caused by heating of the sample (after 400°C, 30 min).

The remarkable discovered strains sustained in the bulk DAP samples undoubtedly modify the biomineral’s properties. As the introduction of compressive residual strains is a common engineering technique to improve material’s mechanical properties (such as tempered glass^38^ or prestressed concrete^39^), such compressive strains present in the DAPs most probably enhance its mechanical properties as well, strengthening the animal’s skeletal integrity. The brittle star’s lenses fulfill an optical function as well, as they guide and focus the light onto photoreceptors inside the tissue.^26^ The presence of high-Mg nanoparticles and associated strains most likely also modify their optical abilities. The presence of well-arranged high-Mg particles^27,29^ may render the crystal photonic, while the discovered mechanical strains associated with the particle layered structure certainly modify the refractive indices of the crystal.^40^ Calcite is birefringent and possesses two different refractive indices: the ordinary one, governing light propagation parallel to its optical axis (*c*-axis), that is larger than the extraordinary one, governing light propagation perpendicular to the optical axis. Since calcite’s refractive indices increase with compressive strains,^40^ and the discovered strains are larger along the *a*-axis than along the *c*-axis, the extraordinary refractive index most probably increases more than the ordinary one, thus reducing the difference between them and diminishing the crystal’s birefringence. Such high anisotropic strains can therefore be advantageous, for the crystal’s optical and mechanical functions in the organism.

## Conclusion

Although the lenses of *O. wendtii* brittle star are a well-studied biomineral, the analysis of XRDT data from bulk DAP sample provided previously unavailable information. These data indeed bear crystallographic information of the large bulk biomineral samples, incorrupt of sample preparation aftereffects. These data showed that the bulk lens possesses two strained states: a stressed, compressed state and a macroscopically unstrained state. Tomographic reconstructions of the XRDT data and comparison with high resolution Nano-HT reconstructions revealed that the undamaged bulk lens was highly strained whereas damaged lens (debris, at cracks, or dents) was macroscopically unstrained. Therefore, mechanical damage induces strain release whose extent can be comprehended by the high intensity difference between the strained and unstrained reflections in those places. The release of strains by cracking manifests an existence of macro-strains caused by gradients of the nanoparticles volume fractions present in the layered structure of Mg-calcite. The unstrained state contains coherent Mg-rich nanoparticles so that Mg-low matrix is still compressed while the particles undergo coherent tensile strains. Moreover, they are such that heating the DAP sample and releasing those strains causes the lenses to burst and collapse via multiple microcracking during coherency loss. In addition to deepening our understanding in the fascinating lenses’ intricate structure, this study highlights the usefulness of XRDT technique, especially in the case of studying biominerals, which can be large, hierarchical, and nanostructured crystals. This study moreover demonstrates the advantage of simultaneously measuring a bulk intact sample with its powder-like counterpart for calibration and data interpretation.

## Methods

### Data acquisition

XRDT was performed at the ID-16B beamline of the ESRF. Bleached samples of DAP were sectioned with a scalpel, and fragments containing lenses were mounted on the tip of a metal needle and fixed with epoxy, and finally placed on a rotating stage. The heated sample was beforehand placed in an oven, at 400°C, for 30mn. For the untreated sample, a 150 nm beam was used, the sample rotated with 1.5° angular steps until 180° was completed. At each of the 120 rotation angles, horizontal line scans with 300 measurements (translation steps of 150 nm) were performed. The data set therefore consists of a 36,000-frame stack: 300 frames at each of the 120 rotation angles. For the heated sample, a 500 nm beam was used, the sample rotated with 2.7° angular steps until 120° was completed. At each of the 44 rotation angles, horizontal line scans with 266 measurements were performed. The energy of the beam was 29.6 keV.

### Data analysis: filter creation

A function that takes as input the raw data and filter it for background and debris was created using the Python environment. For each pixel, and for each rotation angle, the function counts the number of frames (out of the 300 frames at each angle) in which the intensity of the pixel is higher than a certain threshold value (500). When this number of frames is lower than a certain value (50), the function replaces the pixel intensity by a constant value (100, value close to the background value), in all the 300 frames corresponding to the rotation angle. As a result, the intensities generated by the diffraction of debris (which are small and therefore generate intensity for a small amount of translations) and the intensity of the background are replaced by a background value (Supplementary Figure 4).

### Data analysis: tomographic reconstructions

The pixels of interest were chosen on the average frame, by drawing an ROI, using the ImageJ ROI tool. Using this tool, the mean intensity in the ROI was measured through all the 36,000 frames. A sinogram was then created. Before being reconstructed, a background value was subtracted and logarithmic was applied to the sinogram. The processed sinogram was then used for reconstruction, using the TomoJ plug-in.^41^ The SIRT (Simultaneous Iterative Reconstruction Technique) reconstruction algorithm was used, with 250 and 0.2 as iterations number and relaxation coefficient respectively.

### Data analysis: indexing and strains calculations

The sample-detector length (L) was calculated using the {104} bright polycrystalline-like ring, and the corresponding *d*-spacing value (calculated using the lattice parameter extracted from powder HRXRD^27^). Other radii (R) corresponding to different *d*-spacings were calculated using this fitted value, and the relation 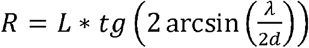. The radial profiles in Figure 5, Figure 6, Supplementary Figure 6 and Supplementary Figure 9 were obtained, with ImageJ, using the plot profile analysis tool. The *x*-axis, in pixels, was converted to 1/*d* units using the previous relation. The differences between the strained and unstrained peaks were determined by fitting the radial profile to a function consisting of the sum of two Gaussian functions (Supplementary Figure 9). The error in Å^−1^ corresponds to a ±0.5-pixel uncertainty value.

## Supplementary material

**Supplementary Figure 1.**
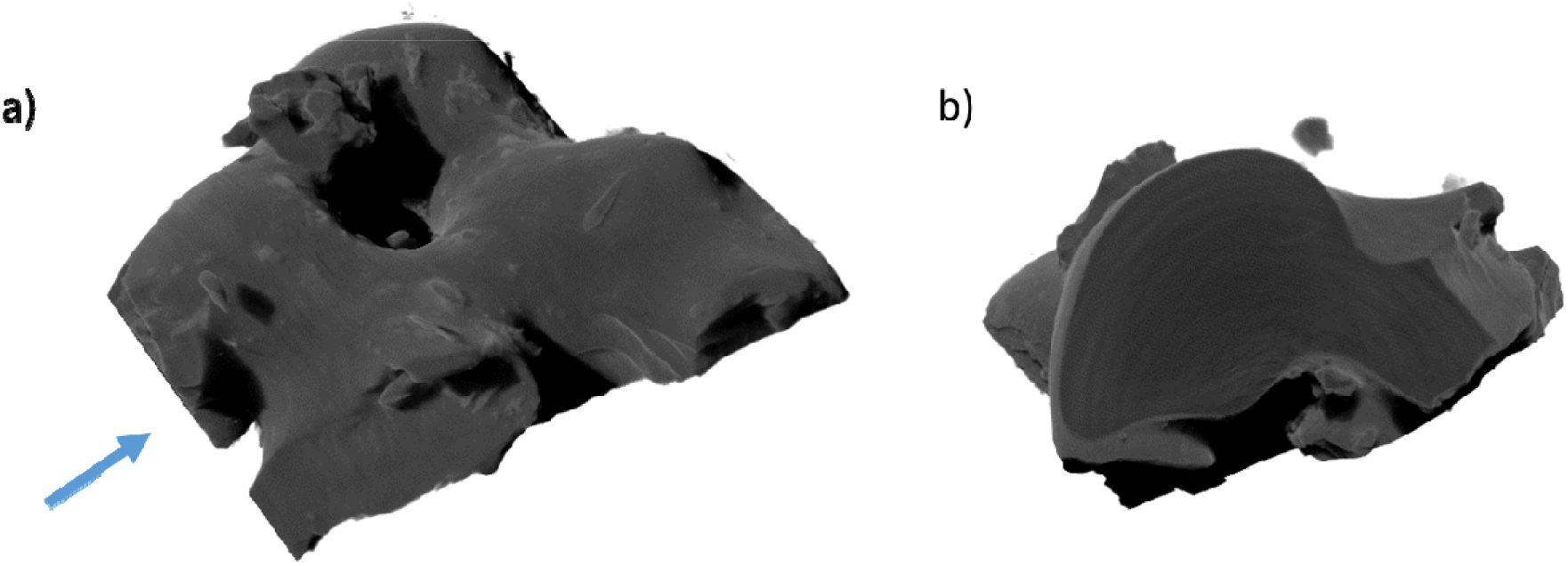
Reconstructed 3D volume rendering of the DAP sample, a) and b) are the same volume from 2 different directions. a) shows the DAP sample is constituted of 3 lenses and debris, the arrow indicates the view direction in b). b) shows the slice measured by XRDT.

**Supplementary Figure 2.**
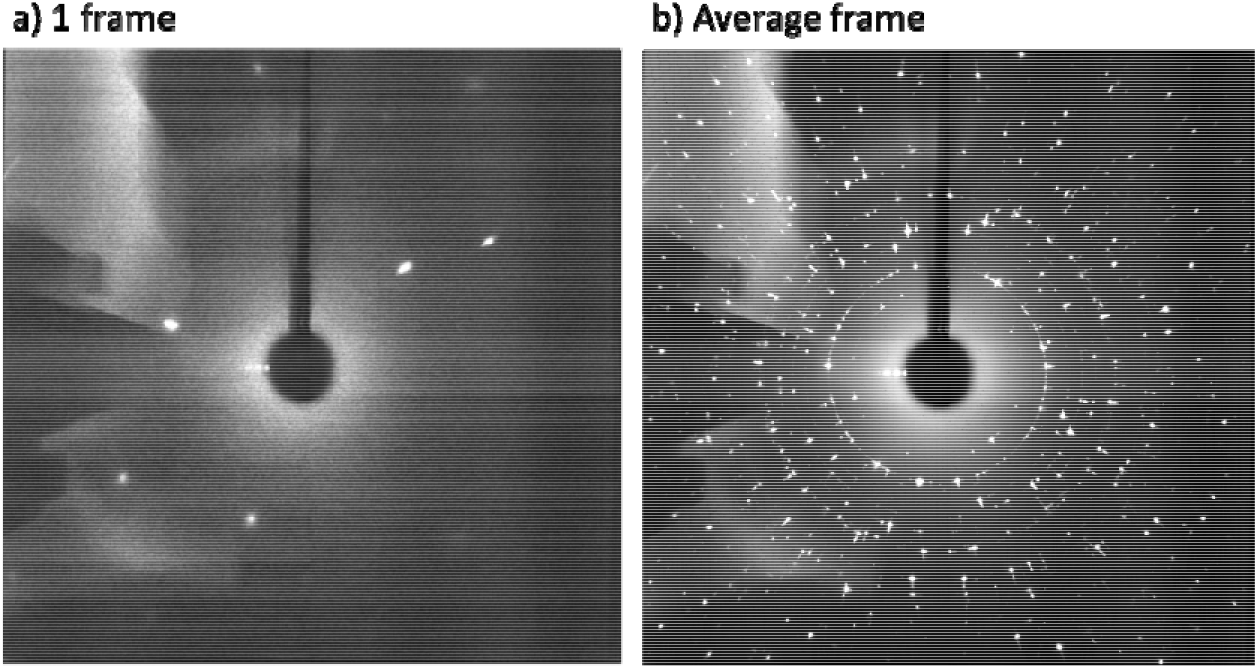
a) 1 frame from the 36,000 data frame stack, b) average frame created with all the frames of the stack.

**Supplementary Figure 3.**
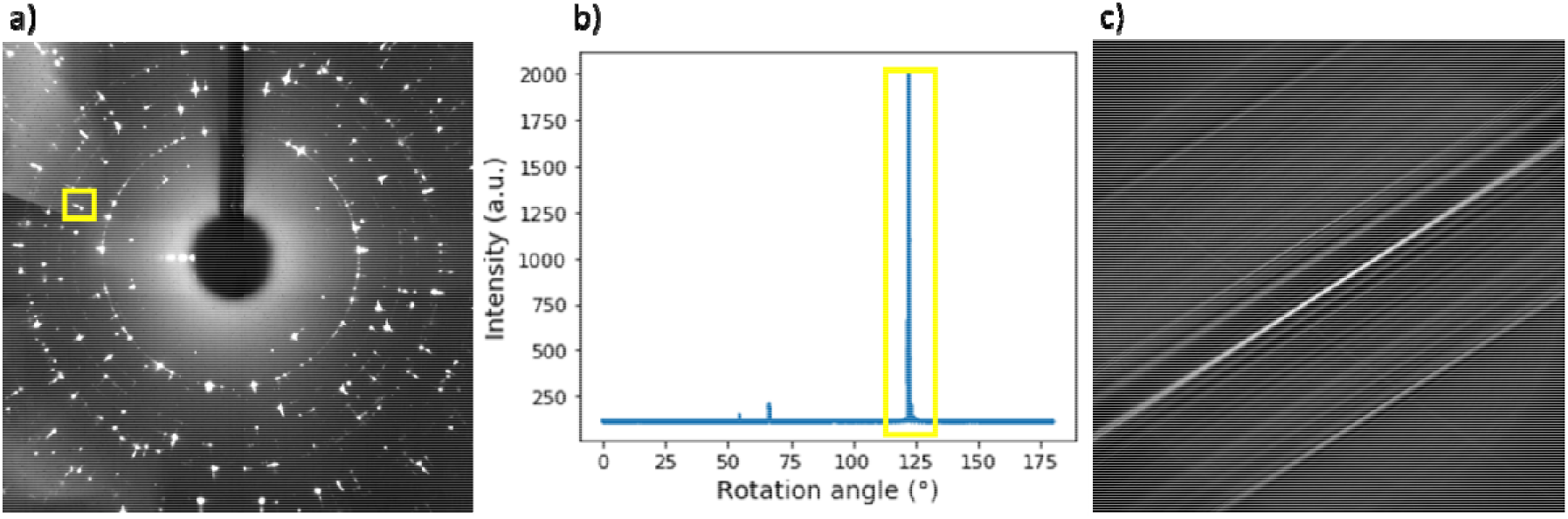
a) Average XRDT frame and the reflection whose pixels were selected as indicated by a yellow rectangle. b) Intensity of the selected pixels through the 36,000 frames showing that the reflection indeed appear at a specific angle only (the 2 small visible peaks correspond to intensity coming from nearby reflections). c) Reconstruction of the selected pixels give information at one angle only (this reconstruction was performed with weighted back projection (WBP) algorithm).

**Supplementary Figure 4.**
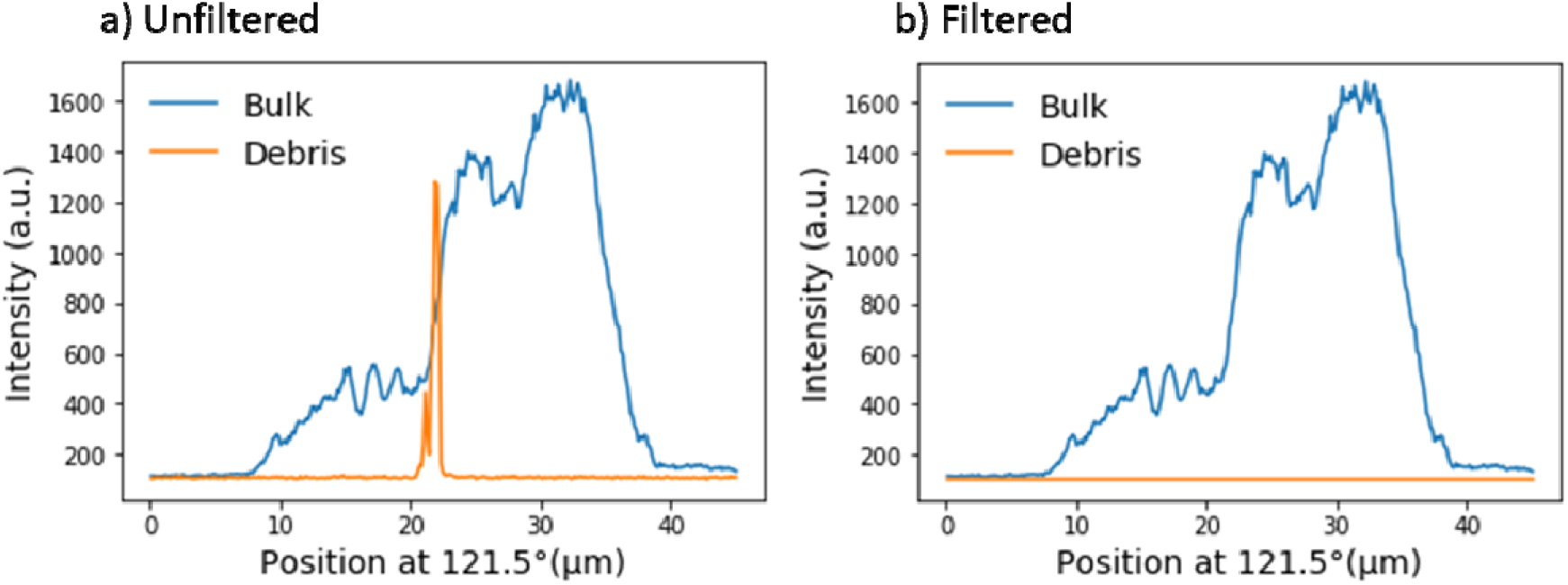
Intensity of pixels belonging to a reflection diffracted by the lens bulk (blue) and by a debris (orange) at a specific angle and through the frames of a) unprocessed data and b) filtered data.

**Supplementary Figure 5.**
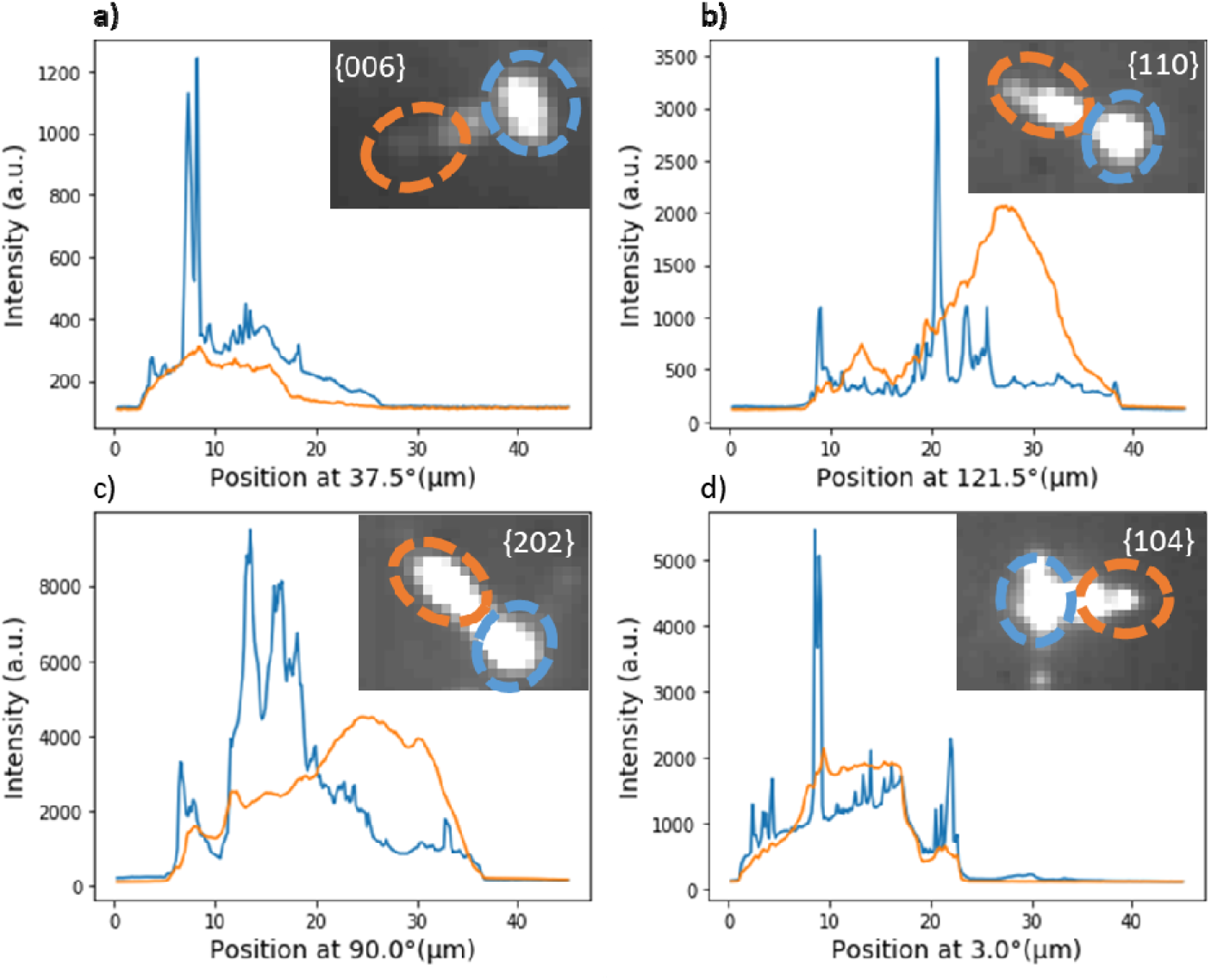
Intensity of unstrained (in blue) and strained (in orange) sides of several reflections. The insets are the considered reflections, the considered regions whose intensities are plotted are marked by dashed circles (unstrained – blue, strained – orange).

**Supplementary Figure 6.**
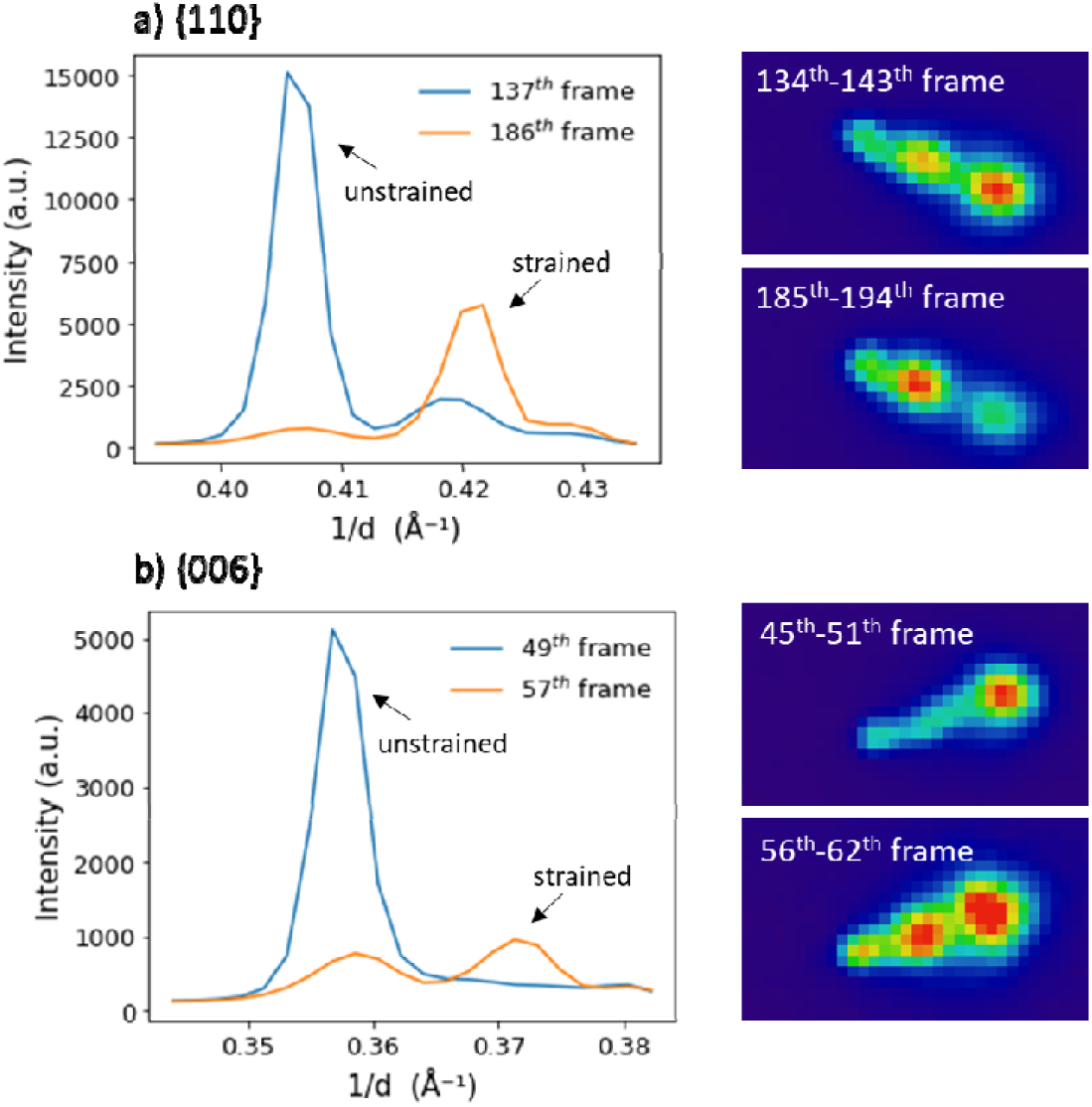
Radial intensity profile of a) {110} and b) {006} reflections in two different frames (of the relevant rotation angle) showing the different relative intensity of the strained and unstrained peak through the frames. On the right are the considered reflections, averaged with neighboring frames.

**Supplementary Figure 7.**
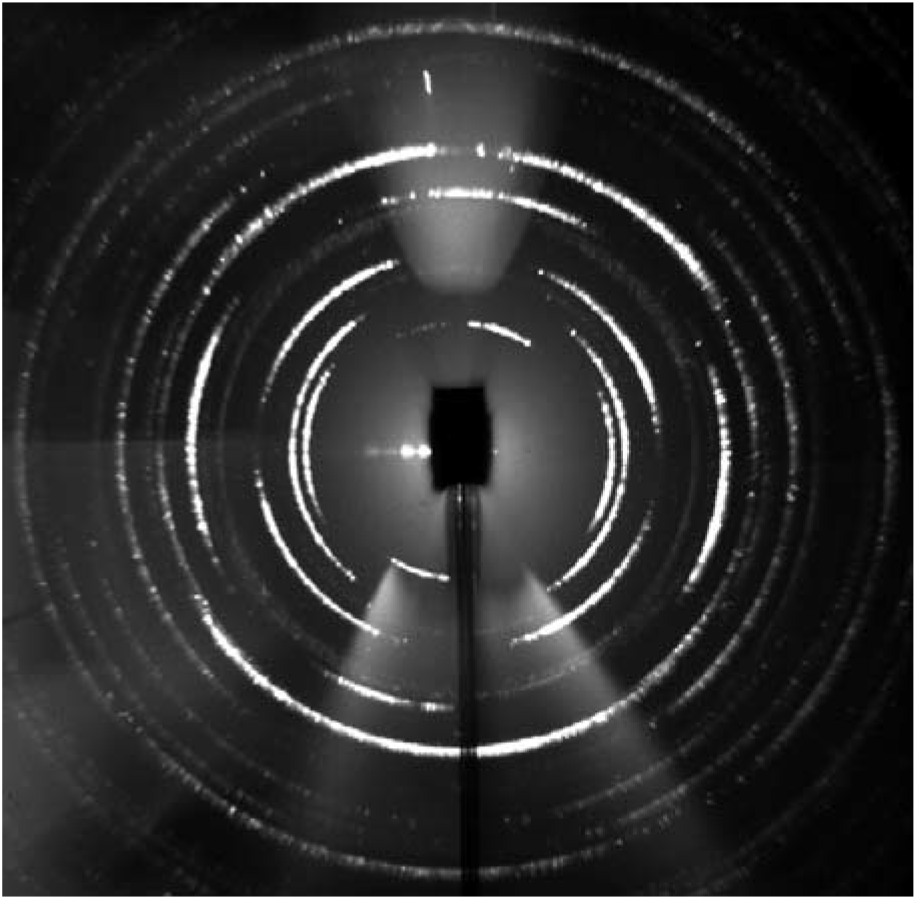
Average frame from an XRDT experiment performed on biogenic vaterite: Herdmania Momus.

**Supplementary Figure 8.**
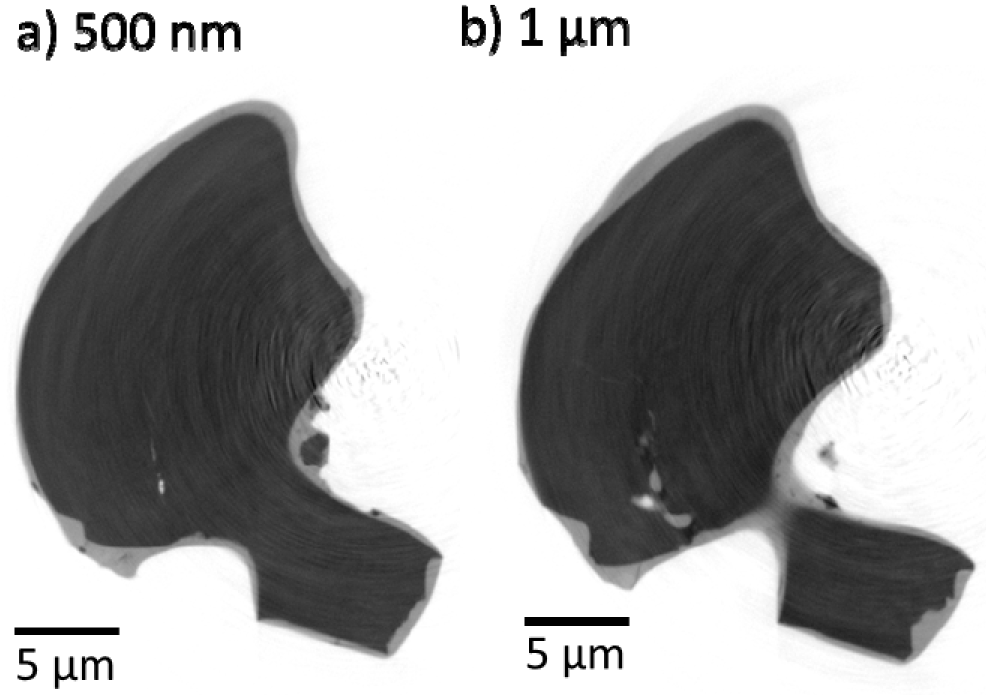
Nano-HT slices of XY planes located at different heights (Z) from the XY plane measured by XRDT: a) 500 nm and b) 1 μm below the plane measured by XRDT, showing the open crack in the lens.

**Supplementary Figure 9.**
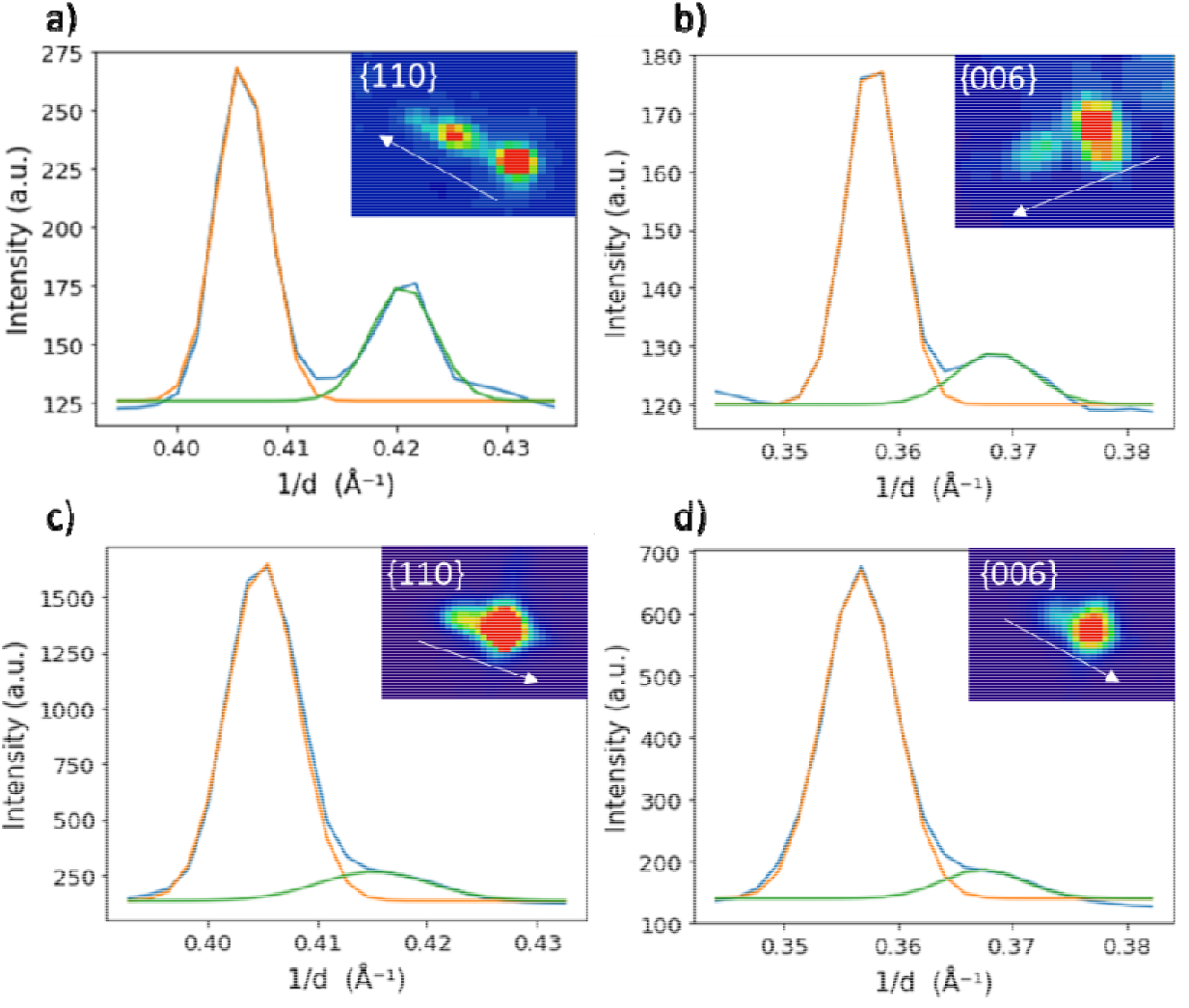
Radial intensity profile and 2-Gaussian fitting of the considered a) {110}, b) {006} reflection of the untreated lens, and c) {110}, d) {006} of the heated lens.

**Supplementary Figure 10.**
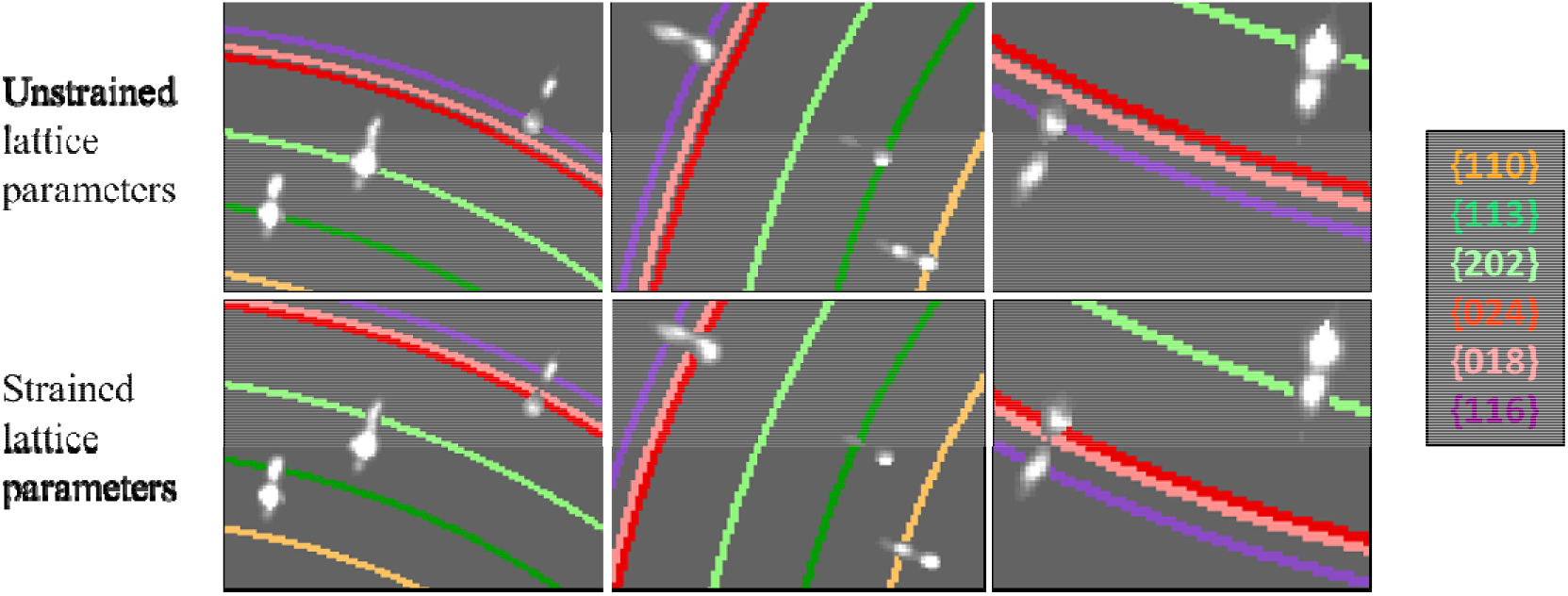
Zoom-ins of the average frame of the filtered data, in which colored circles corresponding to set of planes whose d-spacings are calculated with the unstrained lattice parameters (lattice parameters of powder XRD, top) and strained lattice parameters (bottom) are drawn.

**Supplementary Figure 11.**
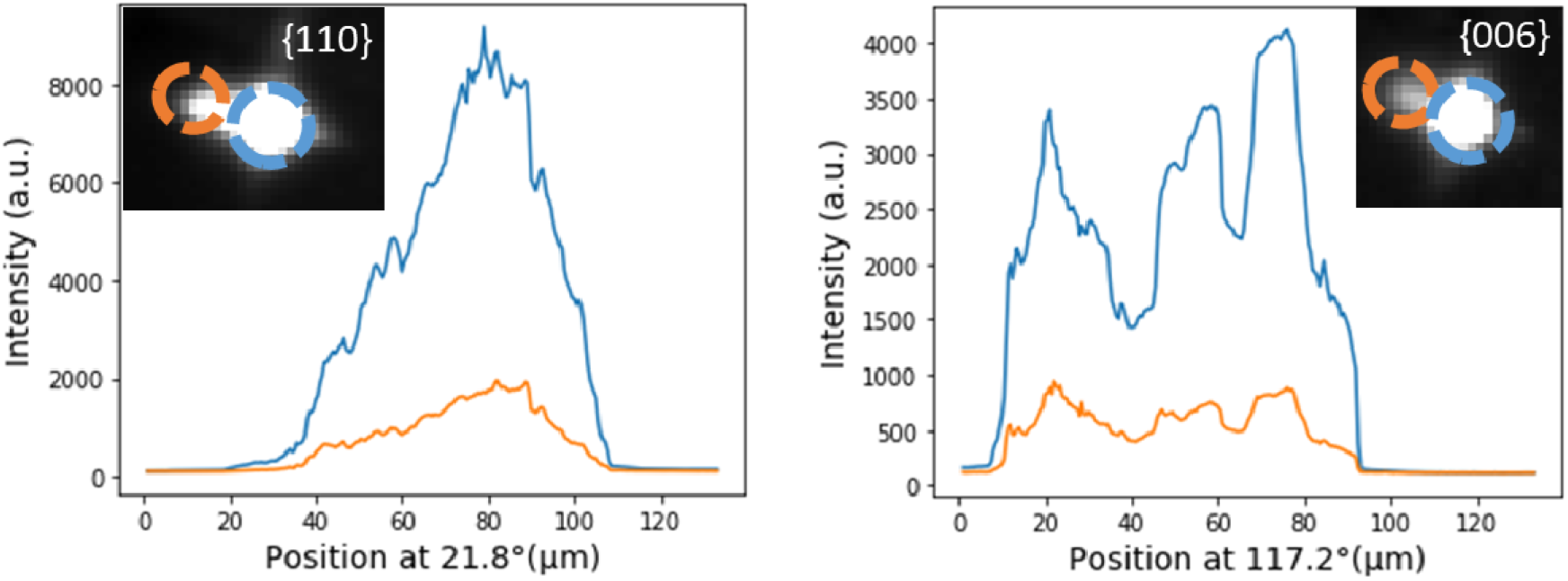
Intensity of unstrained (in blue) and strained (in orange) sides of 2 reflections from the heated sample. The insets are the considered reflections and the considered regions whose intensities are plotted, are marked by dashed circles (unstrained – blue, strained – orange).

